# Aurora B–INCENP localization at centromeres/inner kinetochores is essential for chromosome bi-orientation in budding yeast

**DOI:** 10.1101/515924

**Authors:** Luis J. García-Rodríguez, Taciana Kasciukovic, Viola Denninger, Tomoyuki U. Tanaka

## Abstract

To promote chromosome bi-orientation, Aurora B kinase weakens and disrupts aberrant kinetochore–MT interaction. It has long been debated how Aurora B halts this action when bi-orientation is established and tension is applied across sister kinetochores. Pertinent to this debate, it was shown that Bir1 (yeast Survivin), which recruits Ipl1–Sli15 (yeast Aurora B–INCENP) to centromeres, is dispensable for bi-orientation, raising the possibility that Aurora B localization at centromeres is not required for bi-orientation. Here, we show that the COMA inner kinetochore sub-complex physically interacts with Sli15, recruits Ipl1–Sli15 to the inner kinetochore and promotes chromosome bi-orientation, independently of Bir1, in budding yeast. Moreover, using an engineered recruitment of Ipl1–Sli15 to the inner kinetochore when both Bir1 and COMA are defective, we show that localization of Ipl1–Sli15 at centromeres/inner kinetochores is essential for bi-orientation, refuting the above possibility. Our results give important insight into how Aurora B disrupts kinetochore–MT interaction in a tension-dependent manner, to promote chromosome bi-orientation.

## Introduction

For proper chromosome segregation in mitosis, sister kinetochores must interact with microtubules (MTs) from opposite spindle poles prior to chromosome segregation – this state is called chromosome bi-orientation [1, 2]. To establish chromosome bi-orientation, any aberrant kinetochore–MT interaction must be resolved through a process called error correction. Error correction continues until bi-orientation is established and thus tension is applied across sister kinetochores.

To drive error correction, Aurora B kinase plays a central role – it phosphorylates outer kinetochore components, and this weakens and disrupts aberrant kinetochore–MT interactions [3-5]. However, this action of Aurora B must cease when bi-orientation is established and tension is applied across sister kinetochores. It has long been a topic of debate how tension halts the Aurora B action. One possible explanation is that, upon bi-orientation, sister kinetochores are pulled in opposite directions, stretching the outer kinetochores [6, 7] and thus moving Aurora B substrates away from Aurora B-localizing sites at centromeres/inner kinetochores (spatial separation model) [3, 8, 9]. Consistent with this model, Aurora B delocalizes from outer kinetochores when bi-orientation is established in budding yeast [3, 10], outer kinetochore components are dephosphorylated when tension is applied [11, 12], and ectopic Aurora B targeting to the outer kinetochore destabilizes kinetochore–MT interaction during metaphase in mammalian cells [8].

Activation of Aurora B kinase requires the binding of INCENP. In addition, Survivin needs to bind INCENP to recruit Aurora B–INCENP to centromeres/inner kinetochores. In budding yeast, Aurora B, INCENP and Survivin are called Ipl1, Sli15 and Bir1, respectively [9, 13]. The Aurora B spatial separation model predicts that Aurora B localization at centromeres or inner kinetochores is essential for bi-orientation. However, this notion has been challenged by the observation that, when Sli15 lacks its Bir1-binding domain (Sli15 N-terminus), yeast cells are still able to establish bi-orientation and grow almost normally in the absence of Bir1 [14]. This result has raised the possibility that Aurora B–INCENP localization at the centromere/inner kinetochores is dispensable for chromosome bi-orientation. Alternatively, there might be a Bir1-independent mechanism for recruiting Aurora B–INCENP to centromeres/inner kinetochores, but such a mechanism has not been clearly identified [5, 9].

Here we report that, in budding yeast, the COMA (Ctf19/Okp1/Mcm21/Ame1) kinetochore sub-complex physically interacts with Sli15 and recruits Ipl1–Sli15 to the inner kinetochore independently of Bir1 to promote chromosome bi-orientation. Moreover, by engineering Ipl1–Sli15 recruitment to the inner kinetochores, we show that localization of Ipl1–Sli15 at centromeres/inner kinetochores is essential for bi-orientation.

## Results

### COMA facilitates chromosome bi-orientation independently of Bir1 and not through supporting robust peri-centromere cohesion

If there were a Bir1-independent mechanism of recruiting Ipl1–Sli15 to centromeres or inner kinetochores, regulators of such a mechanism might show negative genetic interaction with *bir1* mutants. In fact, in a recent genome-wide genetic interaction analysis in budding yeast, *bir1-17* mutant showed a strong negative genetic interaction with *mcm21*Δ and *ctf19*Δ [15]. Mcm21 and Ctf19 are non-essential components of the inner kinetochore subcomplex COMA. Moreover, an independent study showed that Sli15 localization at kinetochores was partially diminished in COMA component mutants [16]. These results raised the possibility that COMA recruits Ipl1–Sli15 to inner kinetochores independently of Bir1 to promote bi-orientation and maintain cell viability.

Since *bir1* mutations showed a stronger genetic interaction with *mcm21*Δ than with *ctf19*Δ at physiological culture conditions (growth at 20°C and 27°C) [15], we chose to study the function of Mcm21 further. We fused *BIR1* and *MCM21* with the auxin-induced degron tag (*bir1-aid* and *mcm21-aid*) and investigated how this double depletion affected cell growth (Figure 1A). *STU1* is an essential gene and *stu1-aid* was used as a control for viability [17]. The *bir1-aid* suppressed cell growth in the presence of auxin, but not as completely as did *stu1-aid*. The *mcm21-aid* did not suppress growth on its own but, when combined with *bir1-aid*, showed further growth suppression than did *bir1-aid* alone (Figure 1A). A similar result was obtained using *bir1*Δ (*bir1* deletion rather than depletion) combined with *mcm21-aid* (Figure S1A). Thus, combined depletion (or deletion) of Bir1 and Mcm21 showed a synthetic growth defect.

**Figure 1.**
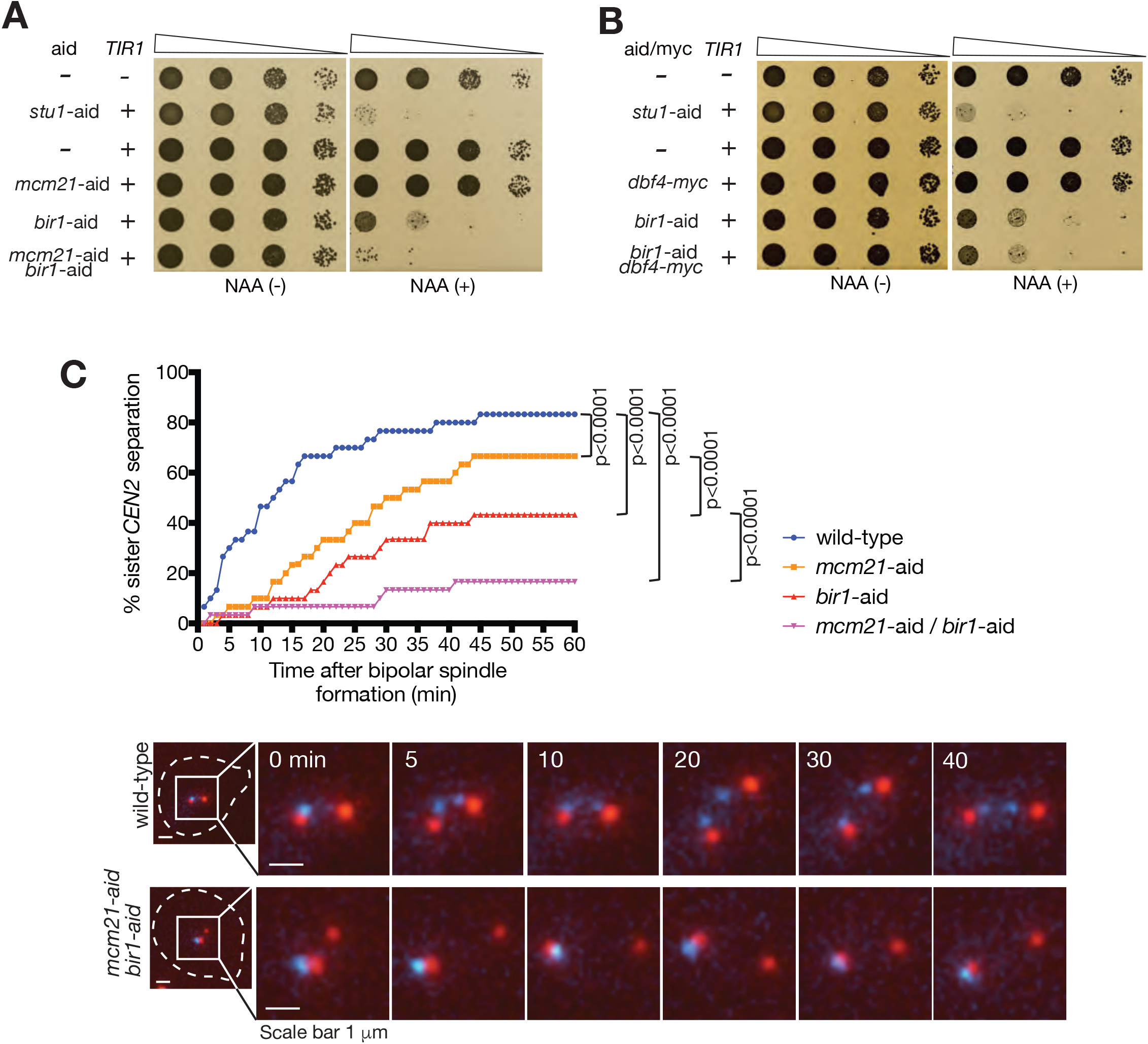
Bir1 and COMA independently promote chromosome bi-orientation. **A.** Bir1 depletion shows synthetic growth defects when combined with Mcm21 depletion. Yeast cells carrying *TIR* with *mcm21-aid* and *bir1-aid* individually and in combination, were serially diluted (10 times dilution each), spotted on plates and incubated for 2 days in the presence (right) and absence (left) of NAA. Cells without AID tag or with *stu1-aid* were analysed in the same way as controls. **B.** Bir1 depletion shows no synthetic growth defects when combined with *dbf4-myc*. Yeast cells carrying *TIR* with *bir1-aid* and *dbf4-myc* individually and in combination were treated and analysed as in A. **C.** Bir1 depletion and Mcm21 depletion cause further defects in chromosome bi-orientation when combined. *MCM21*^*+*^ *BIR1*^*+*^ (wild-type, T12704), *mcm21-aid* (T12697), *bir1-aid* (T12698) and *mcm21-aid bir1-aid* (T12714) cells with *TIR, CEN2-tetOs, TetR-3*×*CFP* and *SPC42-4*×*mCherry* were arrested in G1 with α-factor treatment and released into fresh medium. NAA was added 30 min before the release and also upon release. Microscopy images were acquired from 25 min after the release, for 90 min at 1 min intervals. X-axis shows time relative to separation of spindle pole bodies (Spc42-mCherry), which is defined as time 0. Y-axis shows % of cells showing separation of sister *CEN2s* on the bi-polar spindle (i.e. after SPB separation) for at least two consecutive time points, at or prior to indicated time points. n=30 for each strain; *p*-values were obtained using Kolmogorov–Smirnov test.

The COMA complex promotes robust sister chromatid cohesion at peri-centromere regions by recruiting Dbf4-dependent kinase (DDK) to kinetochores [18-21]. Robust peri-centromere cohesion is important for bi-orientation and cell growth [18, 19, 22]. We addressed whether the above effects of Mcm21 depletion are due to a defect in peri-centromere cohesion. If this is the case, a C-terminus tagged *DBF4* (*dbf4-myc*), which impairs DDK recruitment to kinetochores [20], should show similar behaviour to that of Mcm21 depletion. In fact, *dbf4-myc* expression produced a defect in peri-centromere cohesion in similar extent to (or slightly greater than) that of *mcm21* deletion (Figure S1B). However, in contrast to Mcm21 depletion, *dbf4-myc* showed no synthetic growth defect with Bir1 depletion (Figure 1B). This suggests that the synthetic growth defect shown by combining depletion of Mcm21 and Bir1 is not due to a defect in peri-centromere cohesion.

We compared the efficiency of chromosome bi-orientation establishment in individual Bir1 and Mcm21 depletions with the double depletion. To assay this, we visualized a chosen centromere (*CEN2*) and spindle poles in live-cell fluorescence microscopy, scored the percentage of cells with bi-orientation (separation of sister *CEN2*), and plotted against time after formation of a bipolar spindle (Figure 1C). In a wild-type control, bi-orientation was established in the majority of cells within 15 min. Individual Mcm21 and Bir1 depletions showed moderate and substantial delays, respectively, in bi-orientation establishment. Intriguingly, Mcm21/Bir1 double depletion showed a further delay in bi-orientation than did individual depletions. In the double depletion, only ~17% of cells showed bi-orientation after > 50 min. The effects of Mcm21 and Bir1 depletions seemed to be additive or slightly synergistic, suggesting that COMA and Bir1 independently facilitate chromosome bi-orientation.

Note that, whereas essential COMA components Okp1–Ame1 facilitate outer kinetochore assembly, non-essential components Ctf19–Mcm21 have little such function [23-25]. It is therefore unlikely that the bi-orientation delay in Mcm21 depletion was due to reduced ability of the kinetochore for interacting with MTs. Consistent with this, *CEN2* was always located in the vicinity of one or the other spindle pole (i.e. interacted with MTs) when bi-orientation was defective due to Mcm21 depletion or Mcm21/Bir1 double depletion (Figure 1C, image, bottom).

### COMA physically interacts with Sli15 and recruits Ipl1–Sli15 to the inner kinetochore, independently of Bir1

Independent roles of COMA and Bir1 in facilitating chromosome bi-orientation may be due to their independent functions of recruiting Ipl1–Sli15 to centromeres or inner kinetochores. To address this possibility, we analysed the localization of Ipl1 at centromeres by microscopy. Since centromeres locate on the mitotic spindle, it was difficult to distinguish between Ipl1 localization on centromeres and on spindle MTs. Therefore, to analyse Ipl1 localization specifically at a centromere, we isolated a chosen centromere (*CEN3*) from the spindle by inactivating it (with transcription from the adjacently inserted *GAL* promoter) and thereby disrupting kinetochore–MT interaction [26, 27] (Figure 2A, diagram). Subsequently we shut off transcription from the *GAL* promoter to reactivate *CEN3*, allowing its recapture by spindle MTs (centromere re-activation assay). We analysed Ipl1 localization at *CEN3* after reactivation but before recapture by spindle MTs.

**Figure 2.**
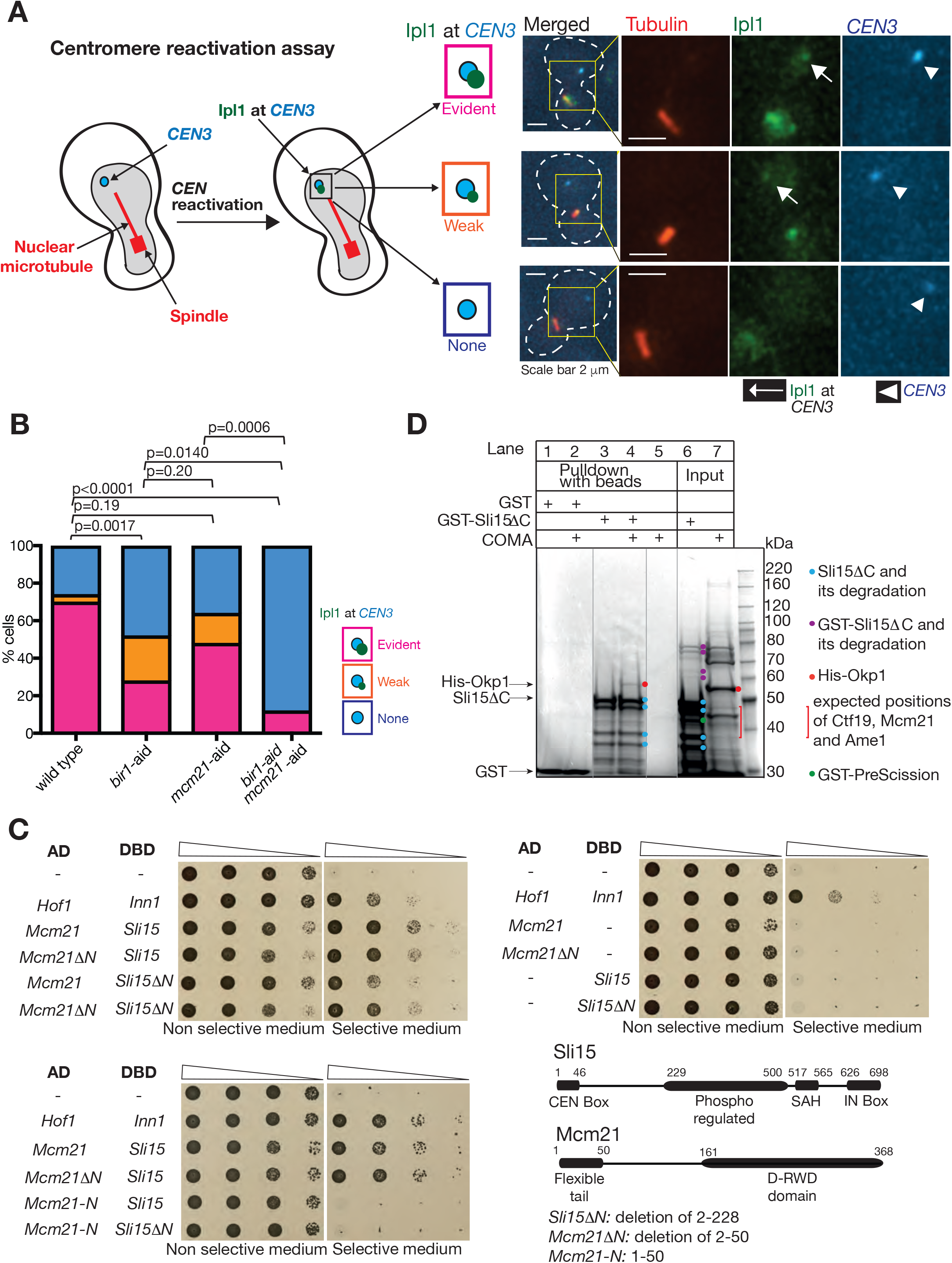
COMA physically interacts with Sli15 and recruits Ipl1–Sli15 to the inner kinetochore independently of Bir1. **A.** Diagram shows the method of analysing Ipl1 signals at isolated *CEN3. CEN3* under *GAL1-10* promoter was inactivated by transcription from the promoter, which prevented interaction with MTs and placed it away from the spindle (diagram, left) [26, 27]. After reactivation of *CEN3* (by shutting off the promoter) during metaphase arrest, Ipl1 at *CEN3* was evaluated (diagram, right). Microscope images show representative examples of no, weak, and evident Ipl1 signals at *CEN3*. **B.** Bir1 and COMA independently recruit Ipl1 to centromeres. *MCM21*^*+*^ *BIR1*^*+*^ (wild-type, T12858), *mcm21-aid* (T12859), *bir1-aid* (T12860) and *mcm21-aid bir1-aid* (T12861) cells with *IPL1-GFP, TIR, GAL1-10* promoter-*CEN3-tetOs, TetR-3*×*CFP, mCherry-TUB1* and *MET3* promoter-*CDC20* were treated and analysed as in diagram in A (see more details in Materials and Methods). Immediately after *CEN3* was reactivated, images were acquired for 10 min with a 1-min interval. Ipl1 signals at *CEN3* were scored into three categories as shown in images in A. n=25–27 for each strain; *p*-values were obtained using chi-squared test for trends. **C.** Mcm21 and Sli15 interact in the yeast two-hybrid assay. The indicated constructs were fused to the Gal4 transcriptional activation domain (AD) and Gal4 DNA-binding domain (DBD). If the AD- and DBD-fused constructs physically interact, yeast cells grow on plates with selective medium. Yeast cells (10 times serial dilution) were incubated at 30 °C for 2 days. Hof1 and Inn1 were used as a positive control [53]. Diagram shows domain structures of Sli15 and Mcm21 [30, 55]. **D.** Recombinant COMA components are pulled down by immobilized recombinant Sli15. The following samples were run on the SDS–PAGE gel and then stained with Coomassie Blue: GST-fused Sli15 (aa 1-401) [GST-Sli15ΔC] and COMA components (including His-tagged Okp1) were expressed in, and purified from, *E. coli* cells (lane 6, 7). GST-Sli15ΔC was immobilized on glutathione magnetic beads; partially purified COMA components were added; washed; and Sli15ΔC was cleaved off from GST by PreScission protease (lane 4). Lane 1, 2, 3 and 5 show controls of the pull-down as indicated. PreScission protease was also added to samples shown in lane 6. Lane 2 and 4 were analysed by mass-spectrometry, and COMA components Mcm21, Ame1 and Okp1, as well as Sli15, were detected in lane 4, but not in lane 2.

Ipl1 localization at *CEN3* was scored in three categories; no localization, weak localization and evident localization (Figure 2A, images). In wild-type control, ~70 % of cells showed evident Ipl1 localization at *CEN3* (Figure 2B). The Ipl1 localization was marginally reduced by Mcm21 depletion and more clearly reduced by Bir1 depletion. Intriguingly, Mcm21/Bir1 double depletion showed greater reduction of Ipl1 localization at *CEN3* compared with that in individual depletions (Figure 2B). The effect of Mcm21 and Bir1 depletions on Ipl1 localization at *CEN3* seemed to be additive or synergistic, suggesting that COMA and Bir1 independently promote Ipl1 localization at centromeres.

We also investigated the effect of *dbf4-myc*, which impairs DDK recruitment to kinetochores (and thus fails to support robust peri-centromere cohesion) [20], on Ipl1 localization on *CEN3* (Figure S2A). *DBF4* wild-type and *dbf4-myc* showed a similar level of Ipl1 localization at *CEN3*. Bir1 depletion reduced Ipl1 localization at *CEN3* but this reduction was similar between *DBF4* wild-type and *dbf4-myc*. These results with *dbf4-myc* contrast with those with Mcm21 depletion (Figure 2B), suggesting that reduced Ipl1 localization at centromeres with Mcm21 depletion in the absence of Bir1 was not due to weakened peri-centromere cohesion.

How then does COMA promote Ipl1 localization at centromeres/inner kinetochores, independently of Bir1? COMA may physically interact with Ipl1–Sli15 to enable their recruitment to the inner kinetochore and we investigated possible physical interactions between Mcm21 and Sli15 using the yeast two-hybrid method (Figure 2C). Indeed, Mcm21 and Sli15 showed physical interaction and this was not dependent on the Bir1-binding domain of Sli15 (Sli15 N-terminus 1–228 aa [28, 29]) or the flexible Mcm21 N-terminus (1–50 aa [30]). We also addressed whether Sli15 directly interacts with COMA, using purified recombinant proteins. It was difficult to purify full-length recombinant Sli15. However, purified and immobilized GST-Sli15 lacking its C-terminus (1–401 aa), pulled down COMA components (Figure 2D). Thus, COMA physically and directly interacts with Sli15, independently of Bir1, to recruit Ipl1–Sli15 to the inner kinetochore.

### Engineered recruitment of Ipl1–Sli15 to the inner kinetochore restores bi-orientation when both COMA and Bir1 are defective

Our results suggest that the level of Ipl1 localization at centromeres is correlated well with efficiency of chromosome bi-orientation when Bir1 and Mcm21 were depleted individually and in combination (Figure 1C and 2B). In particular, with Bir1/Mcm21 double depletion, both Ipl1 localization at centromeres and bi-orientation were almost completely abolished. This raises the possibility that Ipl1–Sli15 localization at centromeres/inner kinetochores is essential for chromosome bi-orientation. One way to test this is to engineer recruitment of Ipl1–Sli15 to centromeres/inner kinetochores in the absence of Bir1 and Mcm21, and to test whether this can rescue bi-orientation.

To engineer Ipl1–Sli15 recruitment to inner kinetochores, we used the rapamycin-dependent association between FRB and FKBP12 [31], and fused FRB and FKBP12 to Sli15 and Mif2 respectively. Mif2 is the yeast CENP-C orthologue and an inner kinetochore component, and was chosen for this purpose since its inner kinetochore localization would not be affected by Mcm21 depletion [25]. As in Figure 2A, we isolated *CEN3* from the mitotic spindle and studied Sli15-FRB localization at *CEN3* in the presence of rapamycin (Figure 3A). When Mif2 was not fused to FKBP12 (control), Bir1 depletion considerably reduced the level of Sli15-FRB at *CEN3*, which is consistent with Figure 2B. Fusion of FKBP12 to Mif2 rescued Sli15-FRB localization at *CEN3* to a normal level, when Bir1 was depleted (Figure 3A). Thus, Sli15 can indeed be recruited to the inner kinetochore by this engineered system.

**Figure 3.**
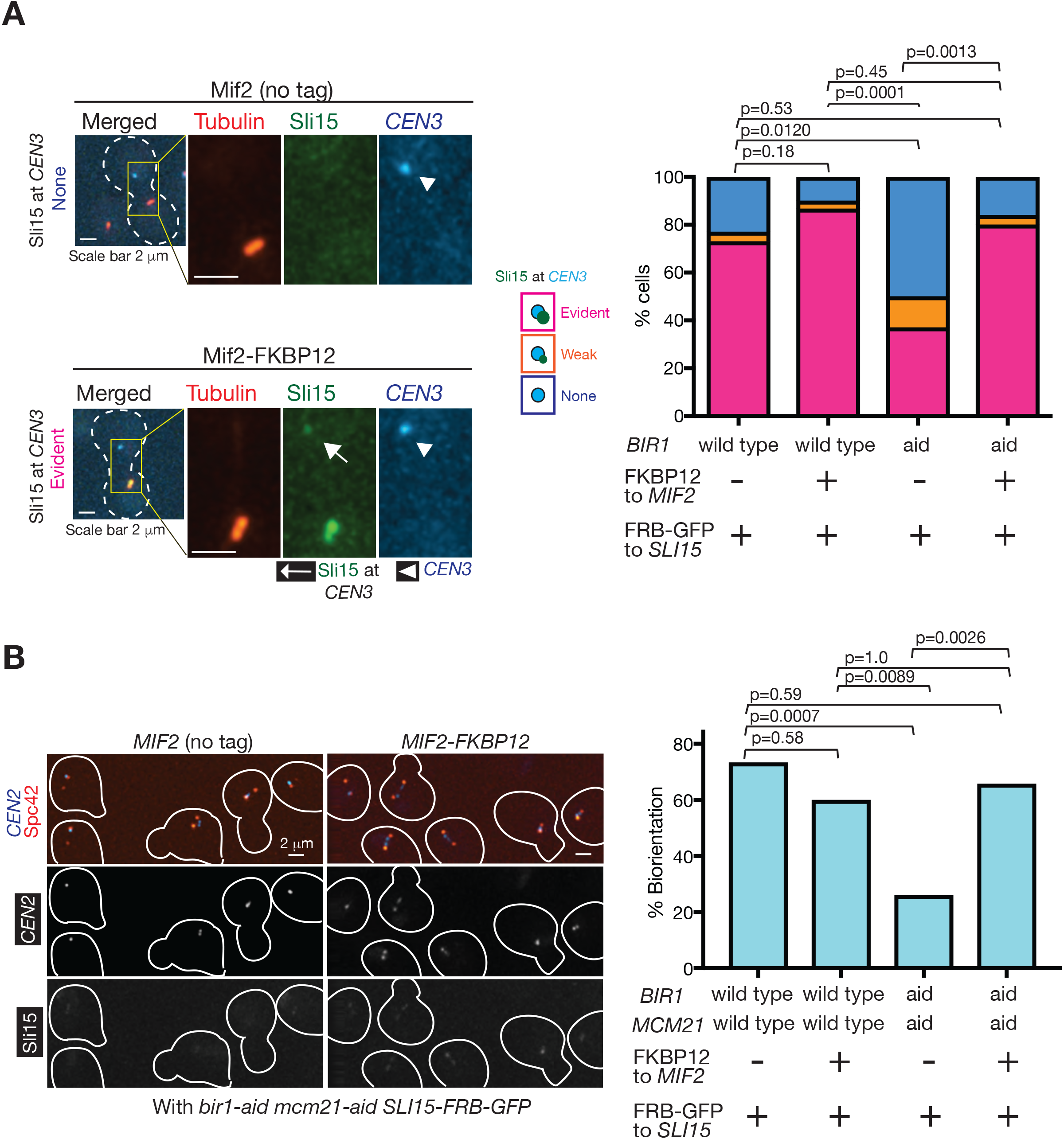
Engineered recruitment of Sli15 to the inner kinetochore restores bi-orientation when both COMA and Bir1 are defective. **A.** FKBP12-fused Mif2 recruits FRB-fused Sli15 to isolated *CEN3*. a) *BIR1+* (wild-type) or *bir1-aid* and b) *MIF2* with or without fusion to *FKBP12*, were combined as indicated below the graph (T13199–T13202 from left to right). All strains carried *SLI15-FRB-GFP, TIR, TOR1-1, fpr1Δ, GAL1-10* promoter-*CEN3-tetOs, TetR-3*×*CFP, mCherry-TUB1* and *MET3* promoter-*CDC20*. Cells were treated as in Figure 2B, except that rapamycin was added 30 min before the start of image acquisition. Sli15-FRB-GFP signals at *CEN3* were scored into three categories as in Figure 2B. n=26–30 for each strain. *p*-values were obtained by chi-squared test for trends. **B.** Engineered Sli15 association with Mif2 restores bi-orientation when both COMA and Bir1 are defective. a) *BIR1*^*+*^ *MCM21*^*+*^ (wild-type) or *bir1-aid mcm21-aid* and b) *MIF2* with or without fusion to *FKBP12*, were combined as indicated below the graph (T13438, T13441, T13440 and T13444 from left to right). All strains carried *SLI15-FRB-GFP, TIR, TOR1-1, fpr1Δ, CEN2-tetOs, TetR-3*×*CFP, SPC42-4*×*mCherry* and *MET3* promoter-*CDC20*. They were cultured in methionine drop-out medium, arrested in G1 with α-factor treatment and released into YPAD plus 2 mM methionine, leading to metaphase arrest (due to Cdc20 depletion). At 2 hours following the release, microscopy images were acquired. Rapamycin was added 30 min before the start of image acquisition. Representative images are shown on left; cell shapes are shown in white lines. Y-axis of the graph (right) shows % of sister *CEN2* separation, representing its bi-orientation. n=30–35 for each strain; *p*-values were obtained using Fisher’s exact test.

We next evaluated efficiency of chromosome bi-orientation with the engineered recruitment of Sli15 to inner kinetochores. As in Figure 3A, FRB and FKBP12 were fused to Sli15 and Mif2, respectively, and bi-orientation frequency was evaluated by visualizing *CEN2* on the spindle in the presence of rapamycin (Figure 3B). When Mif2 was not fused to FKBP12 (control), Sli15-FRB did not co-localize with *CEN2*, and Bir1/Mcm21 double depletion significantly reduced frequency of bi-orientation (as also shown in Figure 1C). Importantly, in the Bir1/Mcm21 double depletion, fusion of FKBP12 to Mif2 allowed Sli15-FRB co-localization with *CEN2*, and rescued frequency of bi-orientation to an almost normal level (Figure 3B). Thus, engineered recruitment of Sli15 to inner kinetochores rescued bi-orientation when both Bir1 and COMA were defective. These results suggest that Ipl1–Sli15 localization at centromeres/inner kinetochores is essential for chromosome bi-orientation.

### Ipl1 still localizes at the centromere/inner kinetochore with *bir1*Δ *sli15*ΔN-terminus, and this is dependent on COMA

It was previously reported that when Sli15 lacks its Bir1-binding domain (Sli15 N-terminus, 1– 228 aa), yeast cells still are able to establish bi-orientation and grow almost normally in the absence of Bir1 [14]. Based on this, it was proposed that Ipl1–Sli15 localization at centromeres is dispensable for chromosome bi-orientation [14]. However, given our finding that COMA promotes recruitment of Ipl1–Sli15 to centromeres/inner kinetochores independently of Bir1 (Figure 2B), bi-orientation and cell growth in *bir1*Δ *sli15*ΔN-terminus may be dependent on COMA. Consistent with this, we found that cell growth of *bir1*Δ *sli15*ΔN-terminus cells was severely reduced when combined with Mcm21 depletion (Figure 4A). In contrast, growth of *bir1*Δ *sli15*ΔN-terminus cells was not affected when deletions were combined with *dbf4-myc* (Figure S2B), which showed a defect in peri-centromere cohesion similar to, or slightly greater than, that of Mcm21 deletion (Figure S1B). Therefore, the effect of Mcm21 depletion on the growth of *bir1*Δ *sli15*ΔN-terminus cells was not due to weakened peri-centromere cohesion, which is found with a lack of Mcm21 [18, 19].

**Figure 4.**
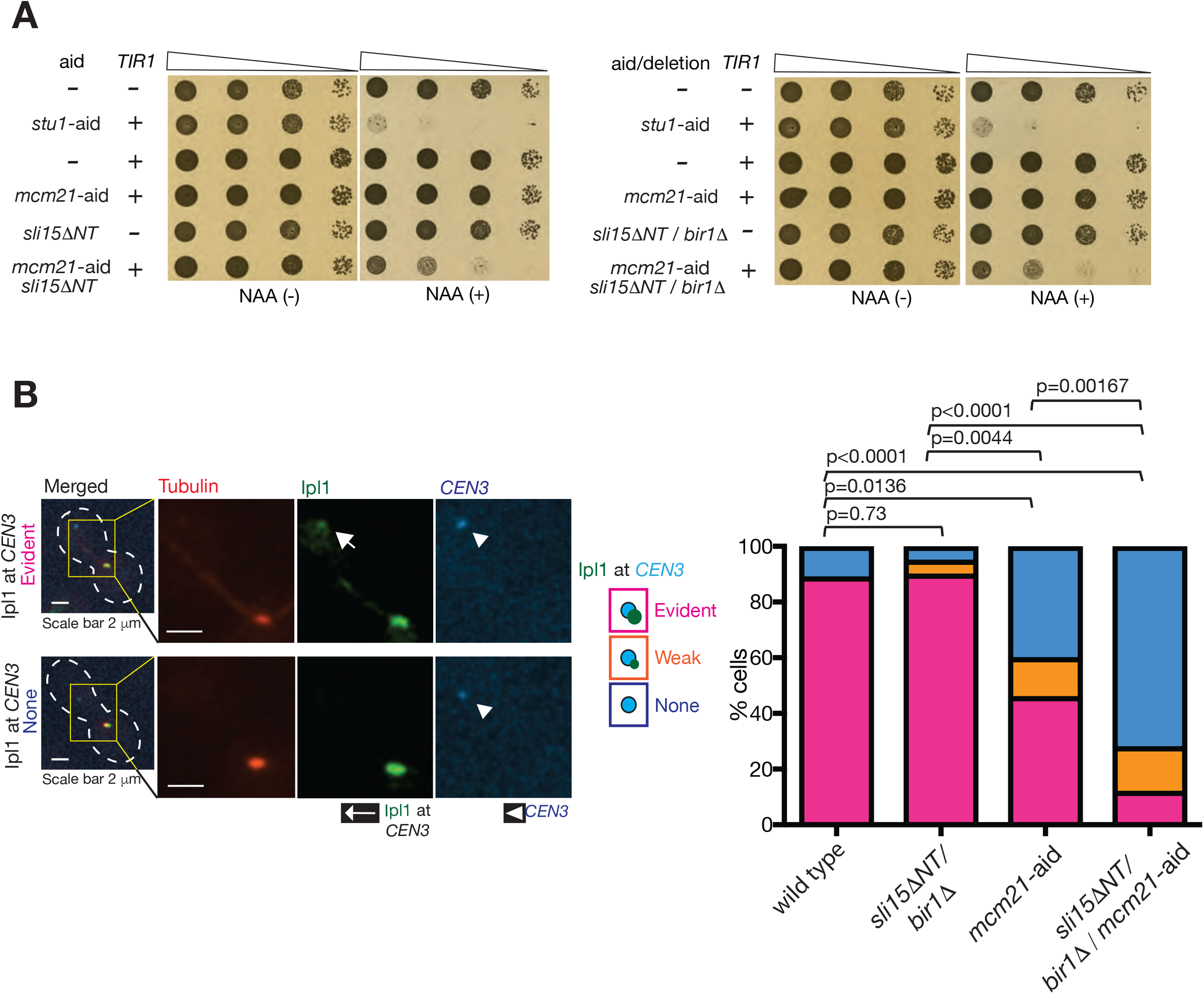
Ipl1 still localizes at the centromere/inner kinetochore with *bir1*Δ *sli15*ΔN-terminus, and this is dependent on COMA. **A.** *bir1*Δ *sli15*ΔN-terminus shows synthetic growth defects when combined with Mcm21 depletion. Yeast cells carrying with *bir1*Δ *sli15*ΔN-terminus and/or *mcm21-aid*, were serially diluted (10 times dilution each), spotted on plates and incubated for 2 days in the presence (right) and absence (left) of NAA. Cells without AID tag or with *stu1-aid* were analysed in the same way, as controls. **B.** Ipl1 localizes at the centromere/inner kinetochore with *bir1*Δ *sli15*ΔN-terminus, dependent on COMA. *BIR1*^*+*^ *SLI15*^*+*^ *MCM21*^*+*^ (wild-type, T122248), *bir1*Δ *sli15*ΔN-terminus (T12229), *mcm21-aid TIR* (T12738) and *bir1*Δ *sli15*ΔN-terminus *mcm21-aid TIR* (T12739) cells with *IPL1-GFP, GAL1-10* promoter-*CEN3-tetOs, TetR-3*×*CFP, mCherry-TUB1* and *MET3* promoter-*CDC20* were treated and analysed as in Figure 2B. Immediately after reactivation of *CEN3*, images were acquired for 10 min with a 1-min interval. Ipl1 signals at *CEN3* were scored into three categories as in Figure 2B. Representative images of T12229 and T12739 cells are shown at top and bottom, respectively. n=15–25 for each strain in graph; *p*-values were obtained using chi-squared test for trends.

In cells with *bir1*Δ *sli15*ΔN-terminus, does Ipl1–Sli15 still localize at centromeres/inner kinetochores and, if so, how is this affected by Mcm21 depletion? To address these questions, we examined localization of Ipl1 on *CEN3*, which was isolated from the spindle as in Figure 2A. Intriguingly, with *bir1*Δ *sli15*ΔN-terminus, Ipl1 localization at *CEN3* was similar to that with the wild-type control (Figure 4B). This contrasts with Ipl1 localization at *CEN3* with Bir1 depletion alone, cells of which showed considerable reduction of Ipl1 there (Figure 2B). This suggests that *sli15*ΔN-terminus enhances Ipl1 localization at centromeres in the absence of Bir1, though the mechanism for this is still unclear. In any case, when combined with Mcm21 depletion, the *bir1*Δ *sli15*ΔN-terminus cells showed a very severe defect in Ipl1 localization at *CEN3* (Figure 4B). In conclusion, Ipl1 still localizes at the centromere/inner kinetochore with *bir1*Δ *sli15*ΔN-terminus, and this localization depends on COMA.

## Discussion

Aurora B kinase (Ipl1 in budding yeast) plays a central role in resolving aberrant kinetochore– MT interactions to promote chromosome bi-orientation. It has been a topic of considerable debate whether Aurora B localization at centromeres/inner kinetochores is essential for chromosome bi-orientation. In this study, we have demonstrated that the COMA kinetochore sub-complex physically interacts with Sli15 and recruits Ipl1–Sli15 to the inner kinetochore, independently of Bir1. Physical interactions between Sli15 and COMA components (Ctf19 and Mcm21) have also been shown in a recently posted preprint, using chemical crosslinking of recombinant proteins [32]. Furthermore, using an engineered system for recruiting Ipl1– Sli15 to the inner kinetochore when both Bir1 and COMA are defective, we were able to show that localization of Ipl1–Sli15 at centromeres/inner kinetochores is essential for bi-orientation in budding yeast. We presume that the centromere/inner kinetochore is a suitable location of Ipl1–Sli15 to enable efficient phosphorylation of outer kinetochores that drives error correction.

Our finding also gives insight into how tension across sister kinetochores halts the Aurora B action of disrupting kinetochore–MT interaction. One popular model for this is the spatial separation model, i.e. tension causes kinetochore stretching, which moves outer kinetochore substrates away from Aurora B-localizing sites at centromeres [3, 8, 9]. This model predicts that Aurora B localization at centromeres is essential for bi-orientation. However, it has been shown that bi-orientation can be established in the absence of Bir1 in budding yeast [14], and this has raised the possibility that Aurora B/Ipl1 localization at centromeres/inner kinetochores is dispensable for bi-orientation – if so, that would rule out the spatial separation model. Our data refute this possibility by demonstrating that the COMA-dependent mechanism still recruits Aurora B/Ipl1 to the inner kinetochore in the absence of Bir1. If our conclusion is correct, the spatial separation model still remains a plausible model for Aurora B-driven error correction, at least in budding yeast.

On the other hand, our data do not exclude other models explaining how tension halts the Aurora B action of disrupting kinetochore–MT interaction. For example, the kinetochore–MT interface may form a stable structure by itself when tension is applied [33], which may overcome the Aurora B action. Localization of Aurora B at the outer kinetochore may have crucial roles in rendering its action tension-dependent [34, 35]. Aurora B kinase activity or counteracting phosphatase activity may be directly regulated by tension [36, 37]. Our finding that Aurora B/Ipl1 localization at the centromere/inner kinetochore is crucial for bi-orientation may also shed new light on these models.

In this study, we used budding yeast as a model organism and concluded that 1) COMA facilitates Ipl1–Sli15 localization to inner kinetochores independently of Bir1, and 2) Ipl1–Sli15 localization at centromeres/inner kinetochores is essential for bi-orientation. Are these also the case in vertebrate cells? Ipl1, Sli15, Bir1 and COMA are conserved from yeast to vertebrates and their vertebrate counterparts are called Aurora B, INCENP, Survivin and CENP-O/P/Q/U, respectively [13, 38]. Crucially, it was recently reported that INCENP lacking its Survivin-binding domain (N-terminus) still supports bi-orientation in human cells [39], as does yeast Sli15 lacking its N-terminus [14]. If mechanisms are conserved between yeast and vertebrate cells, CENP-O/P/Q/R would promote recruitment of Aurora B–INCENP to inner kinetochores independently of Survivin to support error correction in vertebrate cells. Intriguingly, in the *Xenopus* egg extract system, the inner kinetochore was not fully assembled when INCENP lacked its N-terminus (in contrast to human cells); in this circumstance, this INCENP mutant could no longer support error correction even if the outer kinetochore assembly seemed normal [40]. Therefore, inner kinetochore components may indeed support Aurora B–INCENP localization and error correction, independently of Survivin, in vertebrate cells.

## Acknowledgement

We thank Tanaka lab members for discussion; M.J.R. Stark for sharing unpublished results; L. Clayton for editing the manuscript; Dundee Imaging Facility for help in microscopy; Dundee Proteomic Facility for help in mass spectrometry; K. Bloom, R. Ciosk, E.A. Craig, J.E. Haber, M. Kanemaki, K. Labib, U.K. Laemmli, K. Nasmyth. K.E. Sawin, A. Straight, R.Y. Tsien, EUROSCARF and Yeast Resource Centre for reagents. This work was supported by an ERC advanced grant (322682) and the Wellcome Trust (096535 and 097945). T. U. T is a Wellcome Trust Principal Research Fellow.

## Materials and Methods

### Yeast strains and cell culture

The background of yeast strains (W303) and the methods for yeast culture have been described previously [41, 42]. To synchronize cells in the cell cycle, yeast cells were arrested in G1 phase by treatment with yeast mating pheromone (α-factor) and subsequently released to fresh media [41]. Cells were cultured at 25°C in YPA medium containing 2% glucose (YPAD) unless otherwise stated. Constructs of *GAL1-10* promoter-*CEN3-tetOs* [26, 43, 44], *TetR-3*×*CFP* [43, 45], *MET3* promoter-*CDC20* [46], *mCherry-TUB1* [47], *SPC42-4*×*mCherry, NIC96-4*×*mCherry* [48], *CEN2*-*tetO*s (*tetO*×224 was inserted 600 bp away from *CEN2*) [20] and *tetO*s at 15 kb from *CEN12* [20] were described previously. To generate *bir1*Δ and *mcm21*Δ, the whole coding region of the gene was replaced with the *Kl. LEU2* and *KAN-MX4* cassette, respectively, using a one-step PCR procedure [49]. *IPL1* was tagged with yEGFP at its C-terminus at its original locus using the yEGFP-SpHIS5 cassette (pKT128) as a PCR template using a one-step PCR procedure [50]. *sli15*ΔN-terminus (aa 229-698) replaced the original *SLI15+* wild-type gene at its original locus, using the loop-in and loop-out strategy, as follows; 1) *SLI15* promoter, the ATG start codon and *SLI15* coding sequence corresponding to aa 229–465 were cloned into pRS405 vector (URA3 marker) (pT3299), 2) the construct was integrated (loop-in) at *SLI15* locus by homologous recombination after cutting at the *NruI* site within the *SLI15* promoter, 3) the strain was grown with 5-FOA to remove the *URA3* marker (loop-out) by homologous recombination, and 4) strains were checked by PCR and DNA sequencing to select those with *sli15*ΔN-terminus (aa 229–698) and without the original *SLI15*^+^ wild-type gene.

### Depletion of AID-tagged proteins

To deplete Bir1 and Mcm21, *BIR1 and MCM21* were fused to an *aid* tag (auxin-inducible degron tag) at their C-termini at the original loci in the strain carrying the rice F-box gene *TIR1* [51]. In the presence of auxin NAA (1-Naphthaleneacetic acid; 2 mM on plates and 0.5 mM in liquid media), *aid*-tagged proteins interact with the SCF E3 ubiquitin ligase, mediated by Tir1, which leads to their ubiquitylation and degradation by the proteasome [51].

### Engineered association between proteins

To engineer association between Sli15 and Mif2, *SLI15* was tagged with *FRB-GFP* using a *FRB-GFP-kanMX6* (pFA6a-FRB-GFP-kanMX6) cassette, and *MIF2* with *2×FKBP12-TRP1* cassette (pFA6a-2×FKBP12-TRP1) at their C-termini at their original gene loci by a one-step PCR method [52]. For this experiment, yeast strains also carried *TOR1-1*, which conferred rapamycin resistance, and *fpr1*Δ mutations. Association of *FRB* and *FKBP12* fusion proteins was induced by addition of 10 μM of rapamycin to culture media.

### Microscopy image acquisition

During time-lapse imaging, yeast cells were immobilized on a glass-bottomed dish (MatTek, P35G-1.5-10-C) coated with concanavalin A (Sigma C7275), and maintained in synthetic-complete (SC) plus YPA medium (3:1 ratio) [27, 41]. For imaging during metaphase arrest of cells with *MET3* promoter-*CDC20, 2* mM methionine was added to the medium to ensure Cdc20 depletion. Where relevant, NAA was added to the medium during imaging to maintain protein degradation, and Rapamycin was added to the medium during imaging to maintain FRB–FKBP12 interaction. Images were acquired using a DeltaVision Elite microscope (Applied Precision), an UPlanSApo 100× objective lens (Olympus; NA 1.40), SoftWoRx software (Applied Precision), and a CoolSnap HQ (Photometrics). We acquired 7–11 (0.7 μm apart) z-sections, which were subsequently processed through deconvolution, and analysed with Volocity (Improvision) software. CFP, GFP, and mCherry signals were discriminated using the 89006 multi-band filter set (Chroma). For the image panels in Figures, Z sections were projected to two-dimensional images.

### Centromere reactivation assay

To analyse Ipl1 or Sli15 localization at a centromere isolated from the spindle, the centromere re-activation assay was used [26, 27]. In this assay, kinetochore assembly was delayed on a chosen centromere by transcription from the *GAL* promoter (*GAL1-10* promoter-*CEN3-tetOs* replacing *CEN15* on chromosome *XV*). This increased the distance between the centromere and the mitotic spindle, allowing observation of protein localization specifically at *CEN3* after inducing kinetochore assembly on the centromere by turning off the *GAL* promoter in metaphase-arrested cells. Cells with *GAL1-10* promoter-*CEN3-tetOs* and *MET3* promoter-*CDC20* were cultured overnight in methionine drop-out media with 2 % raffinose, treated with α-factor for 2.5 hours (to arrest in G1 phase), and released to fresh media with 2 % raffinose, 2 % galactose and 2 mM methionine (for Cdc20 depletion and *CEN3* inactivation). After 2 hours, cells were suspended in SC medium containing 2 % glucose and methionine to reactivate *CEN3*. Protein localization was analysed at *CEN3* after *CEN3* reactivation and before *CEN3* interaction with MTs extended from the spindle. After background subtraction, Ipl1-GFP signals (colocalising at *CEN3*) of < 5, 5 to 25, and ≥ 25 at their maximum intensity (after Z sections were projected to 2D images) were scored as ‘no’, ‘weak’ and ‘evident’ signals, respectively, using the Voxel Spy tools of Volocity. Sli15-FRB-GFP signals, colocalising at *CEN3*, were scored in the same way, but with those of < 5, 5 to 20, and ≥ 20 as ‘no’, ‘weak’ and ‘evident’ signals, respectively.

### Yeast two-hybrid assay

Two-Hybrid analysis was carried out as in [53]. Briefly, the assay was based on the Gal4 transcription factor and performed after co-transformation of derivatives of pGADT7 (Gal4 activation domain; *LEU2* marker; Clontech) and pGBKT7 (Gal4 DNA binding domain; *TRP1* marker; Clontech) into the yeast strain PJ69-4A (two-hybrid strain [54]). For each assay, independent colonies from the transformation were mixed together in water and spotted in ten-fold dilutions onto SC medium lacking tryptophan and leucine (selective for pGADT7 and pGBKT7, but non-selective for the two-hybrid interaction) and SC medium lacking tryptophan, leucine, histidine and adenine (selective for the two-hybrid interaction).

### Purification of GST-Sli15ΔC

*SLI15* coding DNA, corresponding to aa 1–401, was fused with GST on pGEX6P-1 plasmid (GE Healthcare), which was named pT3176. This construct had a PreScission cleavage site between GST and the SLI15 fragment. The pT3176 was introduced into RosettaGami2 *E. coli* cells (Novagen). The *E. coli* cells were grown in LB medium, and the GST fusion protein was expressed at 18°C with 0.1 mM IPTG induction overnight. Cells were harvested and disrupted using an Emulsiflex cell disruptor (ATA Scientific) in the lysis buffer (50 mM HEPES pH7.6, 1 M NaCl, 5 mM β-mercaptoethanol, 1% Triton X-100) supplemented with cOmplete protease inhibitors (Roche). The cleared lysate was loaded onto 1ml GSTrap FF column (GE Healthcare). Subsequently the proteins, trapped on the column, were eluted with 40 mM reduced glutathione (Sigma) in the elution buffer (50 mM HEPES pH7.6, 0.3 M NaCl, 1 mM DTT, 0.05% NP-40, 10% glycerol). The eluate was loaded onto Superdex 200 10/300 column (GE Healthcare) equilibrated with 50 mM HEPES pH7.6, 0.3 M NaCl, 1 mM DTT, 0.05% NP-40, 15% glycerol, and fractions containing full-length GST-Sli15 (1–401) were pooled and concentrated.

### Partial purification of recombinant COMA

We cloned DNA fragments coding 1) *His*×*6-OKP1* and *AME1* and 2) *CTF19* and *MCM21*, separately into the pETDuet vector (Novagen). The two constructs were introduced to RosettaGami2(DE3)pLysS *E. coli* cells (Novagen), and protein expression was induced with 0.2 mM IPTG at 16°C overnight. Pellets from both cultures were combined, and cells were lysed in buffer 50 mM Tris-HCl pH 7.5, 0.25 M NaCl, 0.5% NP-40, 10 mM β-glycerophosphate, 5% glycerol, 10 mM imidazole and 1 mM PMSF, supplemented with cOmplete protease inhibitors (Roche). The cleared lysate was incubated with His-Pur Nic-NTA Superflow agarose (Thermo). Subsequently the proteins bound on the agarose were eluted in buffer containing 50 mM Tris-HCl pH 7.5, 0.1 M NaCl, 0.1% Triton X-100, 5% 1mM β-mercaptoethanol and 250 mM imidazole. The presence of His-Okp1, Ame1, Ctf19 and Mcm21 in eluates was confirmed by mass spectrometry (MS/MS) after trypsin digestion, and by Western blot with anti-His antibody (Abcam).

### GST pull down assay

GST alone or GST-Sli15 (1-401) was incubated together with partially purified COMA in buffer containing HEPES-NaOH pH 7.4, 150 mM NaCl, 5% glycerol, 0.05% NP-40, 2 mM DTT and 0.5 mM EDTA for 20 min at 4°C. This was added to glutathione magnetic beads (CA-225407, CamBio), which were pre-equilibrated with the above buffer containing 0.1 mg/ml bovine serum albumin. After 1-hour incubation at 4°C with rotation, the beads were washed three times with the above buffer. The proteins, bound to beads via GST, were eluted by two incubations with PreScission protease (GE Healthcare) in the above buffer, and separated by SDS–PAGE (Novex 7% Tris-Acetate gel), and stained with Coomassie Blue (Instant Blue, Expedeon). Note that PreScission protease was also added to the ‘Input’ GST-Sli15 (1-401) sample shown in lane 6 of Figure 2D. We confirmed that the band with the expected size of His×6-Okp1, visible on the Coomassie-stained gel (Figure 2D, lane 4), indeed contained His×6-Okp1 using Western blot with anti-His antibody (Abcam). The areas of the gel, where COMA components were expected to locate, were excised from lane 2 and 4 of Figure 2D, digested with trypsin and analysed by mass spectrometry (MS/MS). COMA components Okp1, Ame1 and Mcm21, as well as Sli15, were present in the PreScission eluate from beads pre-incubated with GST-Sli15 (1-401) and COMA (lane 4), but were absent in the control eluate from beads pre-incubated with GST alone and COMA (lane 2).

---

**Figure S1.**
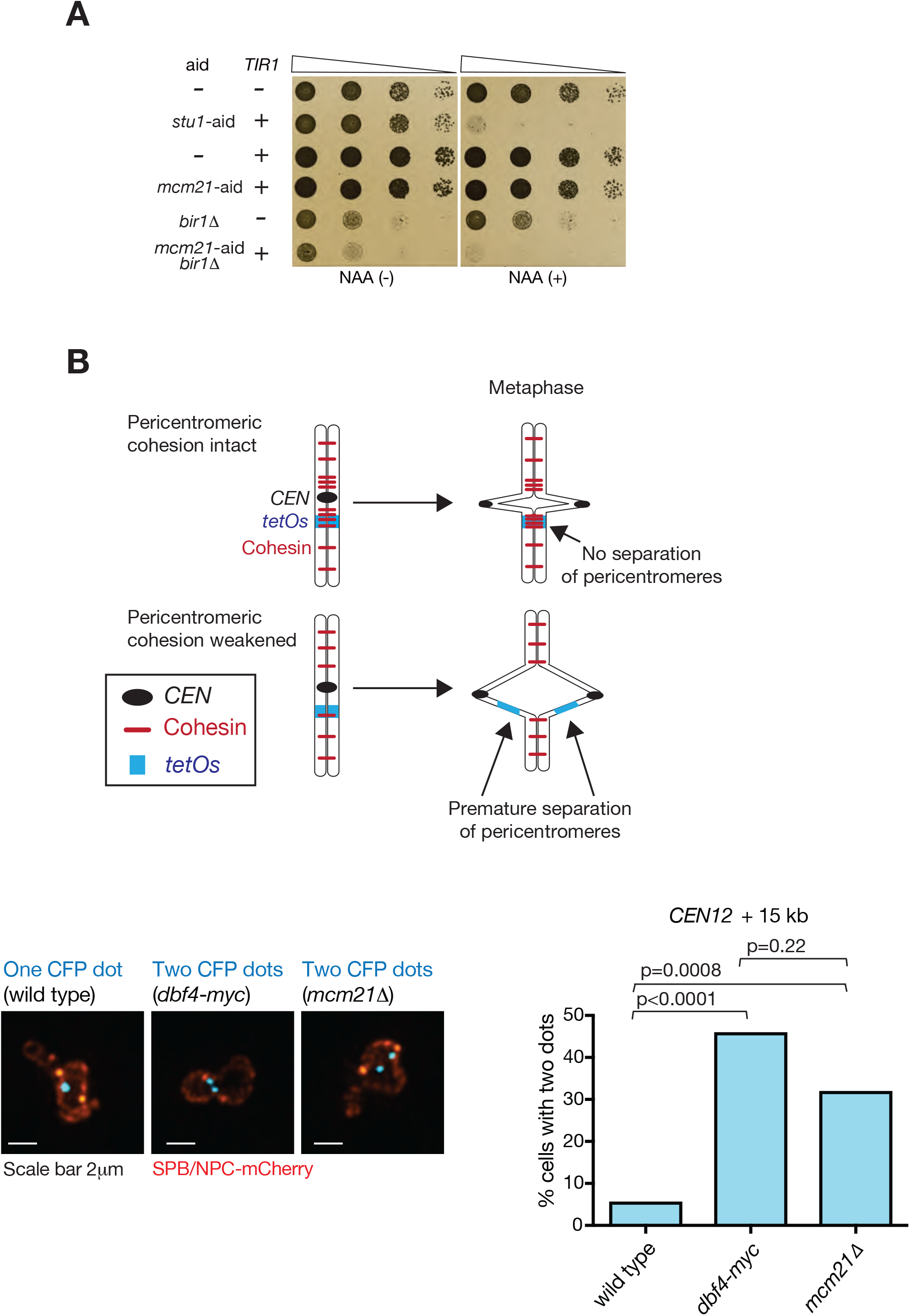
Supplemental figures related to Figure 1. **A.** Bir1 deletion shows synthetic growth defects when combined with Mcm21 depletion. Yeast cells carrying TIR with *mcm21-aid* and *bir1*Δ individually and in combination, were serially diluted (10 times dilution each), spotted on plates and incubated for 2 days in the presence (right) and absence (left) of NAA. Cells without AID tag or with *stu1-aid* were analysed in the same way as controls. **B.** *dbf4-myc* shows a defect in peri-centromere cohesion to a similar extent to (or marginally greater than) *mcm21* deletion. *DBF4*^*+*^ *MCM21*^*+*^ (wild-type, T10141), *dbf4-myc* (T10142) and *mcm21*Δ (T13365) cells with *tetO*s at 15kb from *CEN12, TetR-3*×*CFP, MET3* promoter-*CDC20, SPC42-4*×*mCherry* and *NIC96-4*×*mCherry* were cultured in methionine drop-out medium, arrested in G1 with α-factor treatment and released into YPAD plus 2 mM methionine, leading to metaphase arrest (due to Cdc20 depletion). At 2 hours after the release, microscopy images were acquired. Spc42 and Nic96 are components of the spindle pole body (SPB) and the nuclear pore complex (NPC), respectively, and SPB signals were much brighter than NPC signals. Diagram explains that separation of sister *tetO*s occurs during metaphase when peri-centromere cohesion is weakened. Images show representative examples where sister *tetO*s are, and are not, separated (two and one CFP dots, respectively). Graphs show % of sister *tetO*s separation during metaphase. n=50–52 for each strain; *p*-values were obtained using Fisher’s exact test.

**Figure S2.**
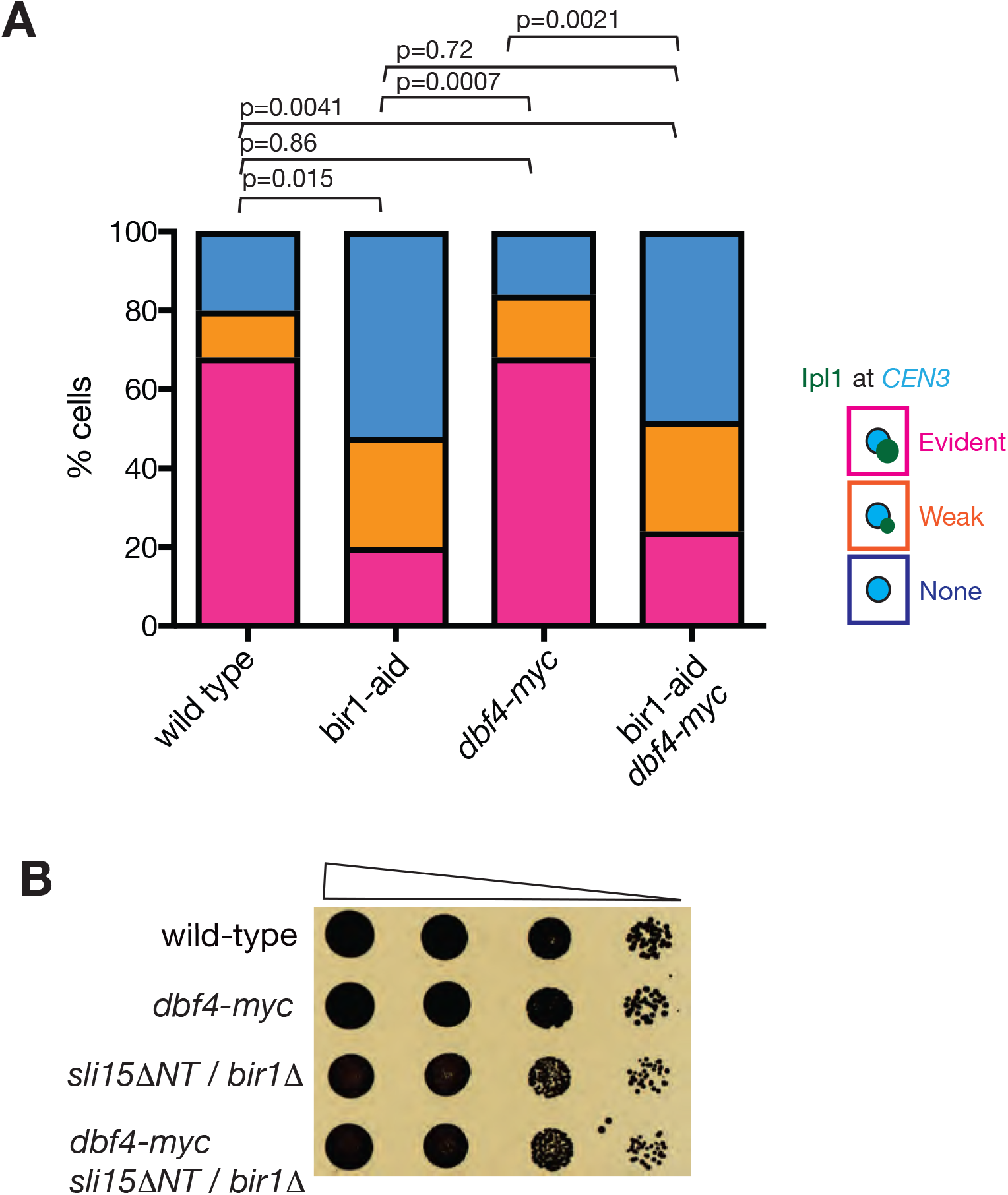
Supplemental figures related to Figure 2 and 4. **A.** *dbf4-myc* does not affect Ipl1 localization at centromeres in the presence and absence of Bir1. *DBF4*^*+*^ *BIR1*^*+*^ (wild-type, T12858), *bir1-aid* (T12860) *dbf4-myc* (T13385) and *bir1-aid dbf4-myc* (T13386) cells with *IPL1-GFP, TIR, GAL1-10* promoter-*CEN3-tetOs, TetR-3*×*CFP, mCherry-TUB1* and *MET3* promoter-*CDC20* were treated and analysed as in Figure 2B. Ipl1 signals at *CEN3* were scored into three categories as in Figure 2B. n=25 for each strain; *p*-values were obtained using chi-squared test for trends. **B.** *dbf4-myc* does not affect growth of cells with *bir1*Δ *sli15*ΔN-terminus. Yeast cells with *bir1*Δ *sli15*ΔN-terminus with *DBF4*^*+*^ or *dbf4-myc*, were serially diluted (10 times dilution each), spotted on plates and incubated for 2 days.

## References

1. Tanaka, T.U. (2010). Kinetochore-microtubule interactions: steps towards bi-orientation. The EMBO journal 29, 4070–4082.

2. Lampson, M.A., and Grishchuk, E.L. (2017). Mechanisms to Avoid and Correct Erroneous Kinetochore-Microtubule Attachments. Biology 6. 197–214.

3. Tanaka, T.U., Rachidi, N., Janke, C., Pereira, G., Galova, M., Schiebel, E., Stark, M.J., and Nasmyth, K. (2002). Evidence that the Ipl1-Sli15 (Aurora kinase-INCENP) complex promotes chromosome bi-orientation by altering kinetochore-spindle pole connections. Cell 108, 317–329.

4. Kelly, A.E., and Funabiki, H. (2009). Correcting aberrant kinetochore microtubule attachments: an Aurora B-centric view. Current opinion in cell biology 21, 51–58.

5. Krenn, V., and Musacchio, A. (2015). The Aurora B Kinase in Chromosome Bi-Orientation and Spindle Checkpoint Signaling. Frontiers in Oncology 5.

6. Uchida, K.S., Takagaki, K., Kumada, K., Hirayama, Y., Noda, T., and Hirota, T. (2009). Kinetochore stretching inactivates the spindle assembly checkpoint. The Journal of cell biology 184, 383–390.

7. Maresca, T.J., and Salmon, E.D. (2009). Intrakinetochore stretch is associated with changes in kinetochore phosphorylation and spindle assembly checkpoint activity. The Journal of cell biology 184, 373–381.

8. Liu, D., Vader, G., Vromans, M.J., Lampson, M.A., and Lens, S.M. (2009). Sensing chromosome bi-orientation by spatial separation of aurora B kinase from kinetochore substrates. Science 323, 1350–1353.

9. Tanaka, T.U., Clayton, L., and Natsume, T. (2013). Three wise centromere functions: see no error, hear no break, speak no delay. EMBO Rep. 14, 1073–1083.

10. Shimogawa, M.M., Widlund, P.O., Riffle, M., Ess, M., and Davis, T.N. (2009). Bir1 is required for the tension checkpoint. Molecular biology of the cell 20, 915–923.

11. Keating, P., Rachidi, N., Tanaka, T.U., and Stark, M.J. (2009). Ipl1-dependent phosphorylation of Dam1 is reduced by tension applied on kinetochores. J Cell Sci 122, 4375–4382.

12. Welburn, J.P., Vleugel, M., Liu, D., Yates, J.R., 3rd, Lampson, M.A., Fukagawa, T., and Cheeseman, I.M. (2010). Aurora B phosphorylates spatially distinct targets to differentially regulate the kinetochore-microtubule interface. Molecular cell 38, 383–392.

13. Biggins, S. (2013). The composition, functions, and regulation of the budding yeast kinetochore. Genetics. 194, 817–846.

14. Campbell, C.S., and Desai, A. (2013). Tension sensing by Aurora B kinase is independent of survivin-based centromere localization. Nature. 497, 118–121.

15. Makrantoni, V., Ciesiolka, A., Lawless, C., Fernius, J., Marston, A., Lydall, D., and Stark, M.J.R. (2017). A Functional Link Between Bir1 and the Saccharomyces cerevisiae Ctf19 Kinetochore Complex Revealed Through Quantitative Fitness Analysis. G3 (Bethesda, Md.) 7, 3203–3215.

16. Knockleby, J., and Vogel, J. (2009). The COMA complex is required for Sli15/INCENP-mediated correction of defective kinetochore attachments. Cell Cycle 8, 2570–2577.

17. Vasileva, V., Gierlinski, M., Yue, Z., O’Reilly, N., Kitamura, E., and Tanaka, T.U. (2017). Molecular mechanisms facilitating the initial kinetochore encounter with spindle microtubules. The Journal of cell biology 216, 1609–1622.

18. Fernius, J., and Marston, A.L. (2009). Establishment of cohesion at the pericentromere by the Ctf19 kinetochore subcomplex and the replication fork-associated factor, Csm3. PLoS genetics 5, e1000629.

19. Ng, T.M., Waples, W.G., Lavoie, B.D., and Biggins, S. (2009). Pericentromeric sister chromatid cohesion promotes kinetochore biorientation. Molecular biology of the cell 20, 3818–3827.

20. Natsume, T., Muller, C.A., Katou, Y., Retkute, R., Gierlinski, M., Araki, H., Blow, J.J., Shirahige, K., Nieduszynski, C.A., and Tanaka, T.U. (2013). Kinetochores coordinate pericentromeric cohesion and early DNA replication by cdc7-dbf4 kinase recruitment. Mol Cell. 50, 661–674.

21. Hinshaw, S.M., Makrantoni, V., Harrison, S.C., and Marston, A.L. (2017). The Kinetochore Receptor for the Cohesin Loading Complex. Cell 171, 72–84.

22. Tanaka, T., Fuchs, J., Loidl, J., and Nasmyth, K. (2000). Cohesin ensures bipolar attachment of microtubules to sister centromeres and resists their precocious separation. Nature cell biology 2, 492–499.

23. Mehta, G.D., Agarwal, M., and Ghosh, S.K. (2014). Functional characterization of kinetochore protein, Ctf19 in meiosis I: an implication of differential impact of Ctf19 on the assembly of mitotic and meiotic kinetochores in Saccharomyces cerevisiae. Mol Microbiol 91, 1179–1199.

24. Hornung, P., Troc, P., Malvezzi, F., Maier, M., Demianova, Z., Zimniak, T., Litos, G., Lampert, F., Schleiffer, A., Brunner, M., et al. (2014). A cooperative mechanism drives budding yeast kinetochore assembly downstream of CENP-A. The Journal of cell biology 206, 509–524.

25. Schmitzberger, F., Richter, M.M., Gordiyenko, Y., Robinson, C.V., Dadlez, M., and Westermann, S. (2017). Molecular basis for inner kinetochore configuration through RWD domain-peptide interactions. The EMBO journal 36, 3458–3482.

26. Tanaka, K., Mukae, N., Dewar, H., van Breugel, M., James, E.K., Prescott, A.R., Antony, C., and Tanaka, T.U. (2005). Molecular mechanisms of kinetochore capture by spindle microtubules. Nature 434, 987–994.

27. Tanaka, K., Kitamura, E., and Tanaka, T.U. (2010). Live-cell analysis of kinetochore-microtubule interaction in budding yeast. Methods 51, 206–213.

28. Jeyaprakash, A.A., Klein, U.R., Lindner, D., Ebert, J., Nigg, E.A., and Conti, E. (2007). Structure of a Survivin-Borealin-INCENP core complex reveals how chromosomal passengers travel together. Cell 131, 271–285.

29. Nakajima, Y., Tyers, R.G., Wong, C.C., Yates, J.R., 3rd, Drubin, D.G., and Barnes, G. (2009). Nbl1p: a Borealin/Dasra/CSC-1-like protein essential for Aurora/Ipl1 complex function and integrity in Saccharomyces cerevisiae. Molecular biology of the cell 20, 1772–1784.

30. Schmitzberger, F., and Harrison, S.C. (2012). RWD domain: a recurring module in kinetochore architecture shown by a Ctf19-Mcm21 complex structure. EMBO Rep 13, 216–222.

31. Chen, J., Zheng, X.F., Brown, E.J., and Schreiber, S.L. (1995). Identification of an 11-kDa FKBP12-rapamycin-binding domain within the 289-kDa FKBP12-rapamycin-associated protein and characterization of a critical serine residue. Proceedings of the National Academy of Sciences of the United States of America 92, 4947–4951.

32. Fischboeck, J., Singh, S., Potocnjak, M., Hagemann, G., Solis, V., Woike, S., Ghodgaonkar, M., Andreani, J., and Herzog, F. (2018). The COMA complex is required for positioning Ipl1 activity proximal to Cse4 nucleosomes in budding yeast. bioRxiv. http://dx.doi.org/10.1101/444570

33. Akiyoshi, B., Sarangapani, K.K., Powers, A.F., Nelson, C.R., Reichow, S.L., Arellano-Santoyo, H., Gonen, T., Ranish, J.A., Asbury, C.L., and Biggins, S. (2010). Tension directly stabilizes reconstituted kinetochore-microtubule attachments. Nature. 468, 576–579.

34. DeLuca, K.F., Lens, S.M., and DeLuca, J.G. (2011). Temporal changes in Hec1 phosphorylation control kinetochore-microtubule attachment stability during mitosis. J Cell Sci 124, 622–634.

35. Caldas, G.V., Deluca, K.F., and Deluca, J.G. (2013). KNL1 facilitates phosphorylation of outer kinetochore proteins by promoting Aurora B kinase activity. The Journal of cell biology 16, 16.

36. Sandall, S., Severin, F., McLeod, I.X., Yates, J.R., 3rd, Oegema, K., Hyman, A., and Desai, A. (2006). A Bir1-Sli15 complex connects centromeres to microtubules and is required to sense kinetochore tension. Cell 127, 1179–1191.

37. Gelens, L., Qian, J., Bollen, M., and Saurin, A.T. (2018). The Importance of Kinase-Phosphatase Integration: Lessons from Mitosis. Trends Cell Biol 28, 6–21.

38. Schleiffer, A., Maier, M., Litos, G., Lampert, F., Hornung, P., Mechtler, K., and Westermann, S. (2012). CENP-T proteins are conserved centromere receptors of the Ndc80 complex. Nature cell biology 14, 604–613.

39. Hengeveld, R.C.C., Vromans, M.J.M., Vleugel, M., Hadders, M.A., and Lens, S.M.A. (2017). Inner centromere localization of the CPC maintains centromere cohesion and allows mitotic checkpoint silencing. Nature communications 8, 15542.

40. Haase, J., Bonner, M.K., Halas, H., and Kelly, A.E. (2017). Distinct Roles of the Chromosomal Passenger Complex in the Detection of and Response to Errors in Kinetochore-Microtubule Attachment. Developmental cell 42, 640–654.

41. Amberg, D.C., Burke, D.J., and Strathern, J.N. (2005). Methods in yeast genetics.

42. Tanaka, K., Kitamura, E., Kitamura, Y., and Tanaka, T.U. (2007). Molecular mechanisms of microtubule-dependent kinetochore transport toward spindle poles. The Journal of cell biology 178, 269–281.

43. Michaelis, C., Ciosk, R., and Nasmyth, K. (1997). Cohesins: chromosomal proteins that prevent premature separation of sister chromatids. Cell 91, 35–45.

44. Hill, A., and Bloom, K. (1987). Genetic manipulation of centromere function. Molecular and cellular biology 7, 2397–2405.

45. Bressan, D.A., Vazquez, J., and Haber, J.E. (2004). Mating type-dependent constraints on the mobility of the left arm of yeast chromosome III. The Journal of cell biology 164, 361–371.

46. Uhlmann, F., Wernic, D., Poupart, M.A., Koonin, E.V., and Nasmyth, K. (2000). Cleavage of cohesin by the CD clan protease separin triggers anaphase in yeast. Cell 103, 375–386.

47. Gandhi, S.R., Gierlinski, M., Mino, A., Tanaka, K., Kitamura, E., Clayton, L., and Tanaka, T.U. (2011). Kinetochore-dependent microtubule rescue ensures their efficient and sustained interaction in early mitosis. Developmental cell 21, 920–933.

48. Li, S., Yue, Z., and Tanaka, T.U. (2017). Smc3 Deacetylation by Hos1 Facilitates Efficient Dissolution of Sister Chromatid Cohesion during Early Anaphase. Molecular cell 68, 605–614.

49. Janke, C., Magiera, M.M., Rathfelder, N., Taxis, C., Reber, S., Maekawa, H., Moreno-Borchart, A., Doenges, G., Schwob, E., Schiebel, E., et al. (2004). A versatile toolbox for PCR-based tagging of yeast genes: new fluorescent proteins, more markers and promoter substitution cassettes. Yeast 21, 947–962.

50. Sheff, M.A., and Thorn, K.S. (2004). Optimized cassettes for fluorescent protein tagging in Saccharomyces cerevisiae. Yeast 21, 661–670.

51. Nishimura, K., Fukagawa, T., Takisawa, H., Kakimoto, T., and Kanemaki, M. (2009). An auxin-based degron system for the rapid depletion of proteins in nonplant cells. Nature methods 6, 917–922.

52. Haruki, H., Nishikawa, J., and Laemmli, U.K. (2008). The anchor-away technique: rapid, conditional establishment of yeast mutant phenotypes. Molecular cell 31, 925–932.

53. Garcia-Rodriguez, L.J., De Piccoli, G., Marchesi, V., Jones, R.C., Edmondson, R.D., and Labib, K. (2015). A conserved Pol binding module in Ctf18-RFC is required for S-phase checkpoint activation downstream of Mec1. Nucleic acids research 43, 8830–8838.

54. James, P., Halladay, J., and Craig, E.A. (1996). Genomic libraries and a host strain designed for highly efficient two-hybrid selection in yeast. Genetics 144, 1425–1436.

55. Fink, S., Turnbull, K., Desai, A., and Campbell, C.S. (2017). An engineered minimal chromosomal passenger complex reveals a role for INCENP/Sli15 spindle association in chromosome biorientation. The Journal of cell biology 216, 911–923.

